# Heterogeneous, population-level drug-tolerant persisters exhibit ion-channel remodeling and ferroptosis susceptibility

**DOI:** 10.1101/2022.02.03.479045

**Authors:** Corey E. Hayford, Blake Baleami, Philip E. Stauffer, B. Bishal Paudel, Aziz Al’Khafaji, Amy Brock, Vito Quaranta, Darren R. Tyson, Leonard A. Harris

## Abstract

Drug-tolerant persisters (DTPs) represent a major obstacle to durable responses in targeted cancer therapy. DTPs are commonly described as distinct single-cell states that survive drug treatment via reversible, non-genetic mechanisms and drive tumor recurrence. Recent work demonstrates that multiple DTPs can coexist, reflecting diversity in lineage, signaling programs, or stress responses. However, each DTP is still generally viewed as a uniform cellular phenotype. Building on our prior work describing a population-level DTP termed “idling” [Paudel et al., *Biophys. J*. (2018) 114, 1499–1511], here we present evidence supporting a fundamentally different view: that DTPs are not single-cell states, but rather heterogeneous populations composed of multiple sub-states with distinct division and death rates that balance to produce near-zero net population growth. Using single-cell transcriptomics and lineage barcoding, we identify multiple phenotypic states within idling DTP populations, with reduced heterogeneity compared to untreated populations, and find that idling DTP cells emerge from nearly all lineages. Transcriptomic and functional analyses further reveal altered ion-channel activity in idling DTPs, which we confirm experimentally. Moreover, drug-response assays reveal increased susceptibility of idling DTPs to ferroptosis, a non-apoptotic form of regulated cell death, indicating the emergence of vulnerabilities associated with drug tolerance. Altogether, our results support a population-level view of tumor drug tolerance in which DTPs comprise stable collections of phenotypic states, shaped by treatment-defined phenotypic landscapes, which are potentially vulnerable to subsequent interventions. This perspective implies that eradicating DTPs will require a fundamental shift away from cell-type-centric strategies toward sequential treatments that progressively reduce phenotypic heterogeneity by modulating the molecular and cellular processes that establish the DTP landscape, an approach previously termed “targeted landscaping.”

## INTRODUCTION

Cancer is a complex, dynamic disease characterized by intratumoral heterogeneity, which has been implicated in treatment evasion and acquired drug resistance (1–3). Tumor heterogeneity is traditionally viewed in terms of clonal cell populations with distinct genetic profiles that pre-exist or emerge during the course of treatment. However, non-genetic sources of tumor heterogeneity are receiving increased attention (4–10). Multiple investigators have reported cancer-cell subpopulations, broadly termed drug-tolerant persisters (DTPs), that withstand drug treatments via non-genetic mechanisms (11–16). DTPs have been variously described as quiescent or slow cycling and as reservoirs from which genetic resistance mutations may emerge. As such, eliminating DTPs before acquired resistance could delay or even prevent tumor recurrence, significantly improving anticancer treatment outcomes. Doing so, however, will require identifying vulnerabilities within the molecular and cellular processes underlying drug tolerance (6, 17–19), which remain poorly understood.

Recently, we described a heterogeneous DTP state, termed “idling,” in BRAF-mutant melanoma cell populations under prolonged BRAF inhibition (17, 20). The idling state is not quiescent but rather characterized by balanced, non-zero rates of division and death, such that the population size remains nearly constant over time. It is also reversible, with the population returning to baseline proliferation after drug removal and reentering the DTP state upon drug reapplication. Mathematical modeling of several drug-tolerant BRAF-mutant melanoma cell populations suggested the idling DTP state is composed of multiple phenotypic sub-states, each with individual rates of division and death, across which cells can transition (17). This characterization of a DTP as a heterogeneous population of single-cell states that contribute collectively to drug tolerance distinguishes idling DTPs from the current consensus view of DTPs (21). Specifically, while prior studies have reported phenotypic heterogeneity within drug-tolerant populations, it has been from the perspective of multiple co-existing DTPs, each considered to be a uniform, single-cell state that either pre-exists or emerges during treatment (11–16). In contrast, while our framework can support the co-existence of multiple DTPs within a population, we add an additional layer of complexity: phenotypic heterogeneity *within* the DTP population itself. This distinction is crucial because cell-type-centric interventions targeting DTPs are likely to fail if DTPs are heterogeneous collections of phenotypic states across which cells can transition during drug treatment. Eliminating any one phenotypic state, or presumed “persister” subtype, will fail to eradicate the drug-tolerant population because cells can reorganize to occupy remaining or newly emergent viable phenotypes. This may explain why effective treatments targeting DTPs have yet to be developed (21), despite numerous ongoing efforts (22–28).

In this work, we use single-cell and bulk RNA sequencing of BRAF-mutant melanoma cell populations to demonstrate that idling DTPs, while less heterogeneous than untreated cell populations, are still composed of multiple phenotypic states. We also use single-cell DNA barcoding to show that idling DTPs emerge from nearly all untreated cell lineages, rather than by competitive selection and expansion of pre-existing drug-tolerant clones. Gene-ontology (GO) analyses and calcium-flux assays indicate that ion-channel activity, including store-operated calcium entry (SOCE), is significantly altered in idling cells. These observations suggest idling DTPs can be induced into ferroptotic cell death (11, 29, 30), which we confirm via drug-response assays demonstrating increased sensitivity to inhibition of glutathione peroxidase 4 (GPX4), a key regulator of ferroptosis. Taken together, we provide experimental support for a population-level view of drug tolerance in which tumor cell populations remodel essential intracellular programs and retain a degree of phenotypic heterogeneity that confounds cell-type-focused targeting. However, vulnerabilities also emerge within DTP populations that may be actionable via secondary or sequential interventions. In what follows, we present the results of our experimental studies probing the transcriptomic, clonal lineage, molecular, and drug-response characteristics of idling DTPs. We conclude with a discussion of the broader implications of the view that DTPs are heterogeneous collections of interchanging phenotypes for future efforts aimed at targeting the biological processes underlying the phenotypic landscape in which these cells reside.

## RESULTS

### Prolonged BRAF inhibition reduces transcriptomic heterogeneity in BRAF-mutant melanoma cell populations

We previously showed that BRAF-mutant melanoma cell populations exhibit complex drug-response dynamics to targeted BRAF inhibition (BRAFi) (17). Across cell lines and single cell-derived subclones, the initial phase of drug response can vary, with proliferation rates ranging from negative to positive. In almost all cases, however, this is followed by entry into a DTP state of near-zero net growth, termed “idling,” in which cell division and death remain balanced. To assess the level of phenotypic heterogeneity in the idling DTP state relative to drug-naïve cells, we performed single-cell RNA sequencing on untreated and idling (8 μM PLX4720 treatment for eight days; see Materials and Methods) BRAF-mutant melanoma cell populations. When the high-dimensional transcriptomic data are visualized using a two-dimensional Uniform Manifold Approximation and Projection (UMAP) (31), we see that the untreated and idling DTP cells reside in distinct regions of the projected space, with minimal overlap (FIG. 1A). We also identify at least two distinct clusters in each case, which we refer to as “large” and “small” untreated and idling DTP clusters (FIGS. 1A and S1A). Note that the large and small untreated clusters are farther apart in UMAP space than the two idling clusters are from each other. Although Euclidian distances in low-dimensional projections must be viewed with caution (31), this result may indicate increased similarity among cells in the two idling clusters relative to cells in the untreated clusters. Taken together, these results suggest that phenotypic heterogeneity is reduced in idling DTPs relative to untreated melanoma cell populations. These observations are consistent with our prior work (17), in which we proposed that a “drug-naïve epigenetic landscape” composed of multiple phenotypic states is modified upon exposure to BRAFi, and that the resulting “drug-modified epigenetic landscape” also comprises multiple phenotypic states but is less heterogeneous overall.

**Figure 1:**
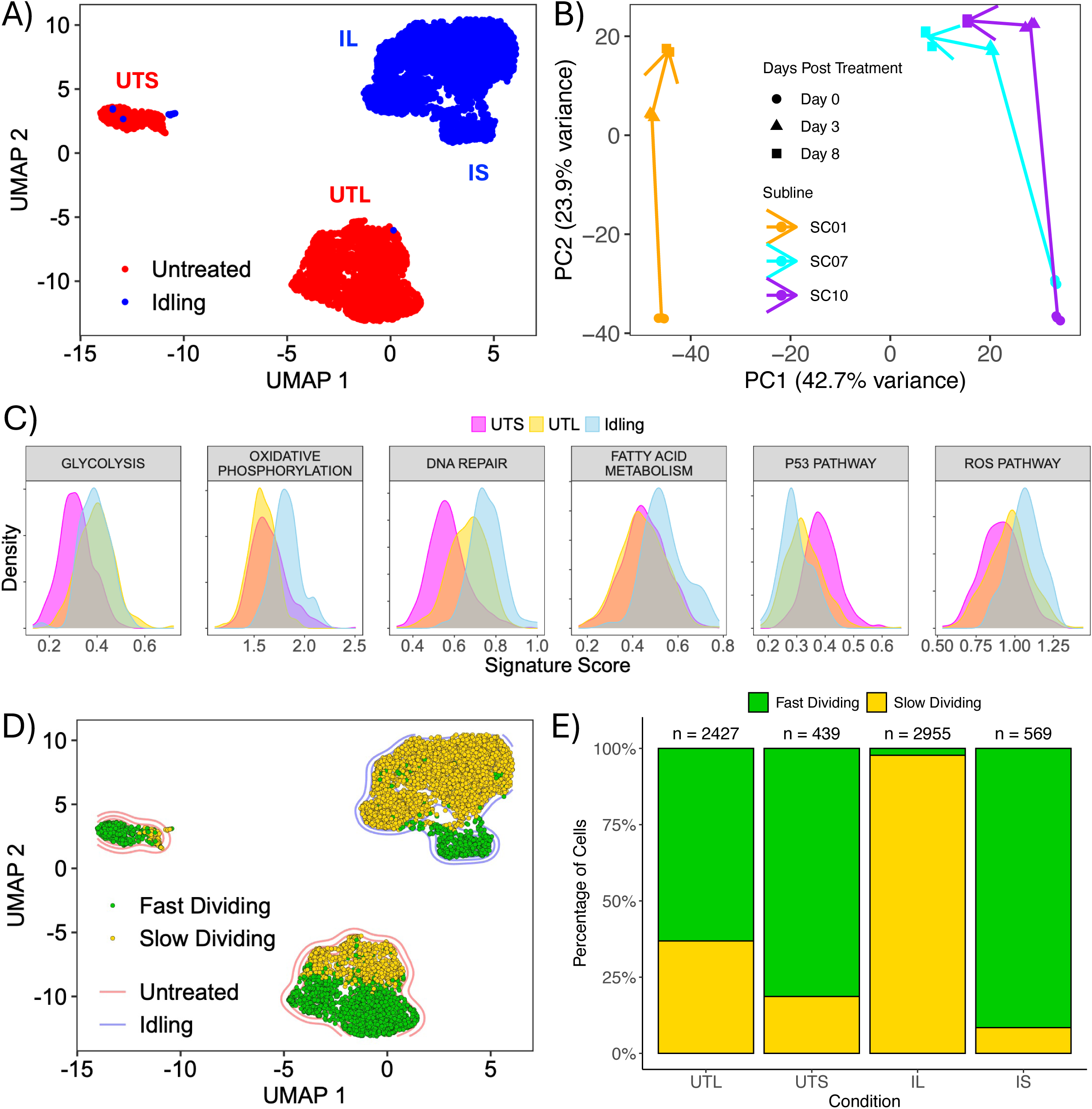
Idling DTPs are composed of a heterogeneous collection of phenotypic states. **(A)** Two-dimensional UMAP space of untreated and idling DTP single-cell transcriptomes. A total of 6410 cells are shown, split ∼50/50 across conditions. UTS: untreated small, UTL: untreated large, IS: idling DTP small, IL: idling DPT large. **(B)** Principal component analysis (PCA) based on bulk RNA sequencing of three drug-treated single-cell-derived subclones (SC01, SC07, SC10) at multiple time points (0, 3, 8 days). All experiments were performed in triplicate. Lines are drawn between the centroids of the triplicates across the time series. **(C)** Distributions of gene signature scores from the functional interpretation analysis for cells in the UTS, UTL, and idling DTP clusters (considered as one cluster in this analysis) in *A*. **(D)** Overlay of cell-cycle state (“fast” = S-G2-M, “slow” = G1) onto all single cells in the single-cell transcriptomic UMAP space. Colored contours (blue and red) behind the cells are cell densities calculated from *A***. (E)** Relative proportions of cells in each cell-cycle state for the two untreated and two idling DTP clusters in *D*. Numbers of cells in each cluster (n) are shown above each bar.

As additional evidence for the reduced phenotypic heterogeneity of the idling DTP state, we performed bulk RNA sequencing on three single-cell-derived subclones of the same BRAF-mutant melanoma cell line (Materials and Methods). These subclones were previously shown to have disparate short-term BRAFi responses, but all transition into an idling DTP state within 3−8 days of treatment (17). Using a principal component analysis (PCA), we see that prior to BRAFi treatment (day 0) there is a significant difference along the first PCA axis (PC1) between one subclone and the other two (FIG. 1B). After BRAFi treatment, all subclones move along the second PCA axis (PC2) and then begin to converge along PC1 by the end of the experiment (day 8), as the populations reach the idling DTP state. Since these subclones are single-cell-derived, it is unlikely these changes are genetic in origin. Therefore, we interpret these results as further evidence that the idling DTP state is less heterogeneous than the untreated population, but still retains a degree of phenotypic heterogeneity.

### Untreated and idling DTP transcriptomic states are associated with different metabolic processes and cell cycle stages

To determine the biological factors that distinguish the single-cell transcriptomic clusters identified in the untreated and idling DTP populations (FIGS. 1A and S1A), we performed a functional interpretation analysis using hallmark gene sets from the Molecular Signatures Database (MSigDB) (32–34) (Materials and Methods). For the untreated population, we considered the large and small clusters separately, across which the population is split approximately 80/20. We see that the large untreated cluster is enriched in a glycolysis gene signature compared to the small untreated cluster (FIG. 1C), which is consistent with our previous report of metabolic heterogeneity within untreated BRAF-mutant melanoma cell populations (20). Furthermore, the large untreated cluster has a higher DNA repair gene signature score, while the small cluster is enriched in a p53 pathway gene signature (FIG. 1C). Comparing the idling population (considered as a single cluster in this analysis) to both untreated clusters, we see enrichment in gene signatures for DNA repair, fatty-acid metabolism, reactive oxygen species (ROS) pathways, and oxidative phosphorylation (FIG. 1C). The latter is consistent with multiple reports in the literature that BRAF-mutant melanoma cells rely primarily on glycolysis for growth and energy production (35, 36) and treatment with BRAFi interferes with glycolytic processes, presumably favoring a switch to oxidative phosphorylation (27, 37).

We also performed differential expression analyses between the two untreated single-cell transcriptomic clusters (FIG. S1B) and between the two idling DTP clusters (the large and small idling clusters are considered separately in this analysis; FIG. S1C). After regressing out cell-cycle-associated genes (along with various other genes; see Materials and Methods), a GO over-enrichment analysis (38) returned numerous cell-cycle-related GO terms (e.g., “nuclear division”, “DNA replication”; FIG. S1C). Thus, even with cell cycle genes removed from the analysis, cell cycle-related processes emerge as major factors distinguishing cells in the large and small idling DTP clusters. To further explore this result, we used a previously established cell cycle gene signature (39) to classify individual cells (both untreated and idling) by cell cycle stage (G1, G2/M, S). For visual clarity, we condensed the cell cycle stages into “slow dividing” (G1) and “fast dividing” (S-G2-M) categories and colored all cells in the single-cell transcriptomics UMAP space by category (FIG. 1D). Within the large and small untreated clusters, we see both slow- and fast-dividing cells, although they are relatively well separated into different regions within the clusters. This may indicate the presence of phenotypic sub-states within the two untreated clusters, which would be consistent with our recent report on transcriptomic heterogeneity within untreated EGFR-mutant non-small cell lung cancer cell populations (6). Conversely, the large idling DTP cluster is highly populated by slow-dividing cells, while the small cluster is composed almost entirely of fast-dividing cells (FIG. 1D,E). Furthermore, the proportion of fast-dividing cells is much lower in the idling DTP population (∼15%) than in the untreated population (∼67%), which is consistent with BRAFi prolonging time spent in G1 (40–42) and the reduced (near-zero) proliferation rate of idling DTPs. Altogether, these results and those from the functional interpretation analysis above (using MSigDB), further support the view that the idling DTP state comprises a heterogeneous collection of phenotypic states.

### Idling DTP cell populations comprise cells from most untreated lineages

To determine whether cells in the idling DTP population are clonally selected from a pre-existing subpopulation or emerge from all untreated lineages after drug treatment, we barcoded the same BRAF-mutant melanoma cell line as above with a guide RNA (gRNA) barcoding library (43, 44). This barcoding system, called ClonMapper (45), establishes a direct connection between the abundance of a clonal lineage and the positions of cells within the single-cell transcriptomics UMAP space (Materials and Methods). Upon treatment with BRAFi for eight days, we see only modest changes in the relative abundances of individual barcodes between the untreated and idling DTP populations (FIGS. 2A and S2A). We also see that the barcode library complexity is reduced by less than 10% (FIG. S2B), barcodes are largely shared across replicates for both the untreated and idling DTP populations (FIG. S2C), and fold-changes in barcode abundances are approximately normally distributed with a mean of zero (FIG. 2B). Notably, nearly all the lineages that do not survive BRAFi treatment come from clones with exceedingly small abundances (data not shown), suggesting that the loss of those lineages is due to random chance (6, 46–48).

Additionally, we mapped barcoded cells for the 24 most abundant barcodes onto the single-cell transcriptomics UMAP space (FIGS. 2C and S2D). In all cases, we see barcoded cells residing in all untreated and idling DTP clusters. Moreover, the changes in the relative abundances of barcodes (FIGS. 2A and S2A) correspond to the proportion of cells in a lineage that reside in the fast-dividing (small) idling DTP cluster, which we assume is populated randomly as the cell population re-equilibrates following drug treatment (17). Lineages in which only a few cells (<5%) reside in the fast-dividing cluster (e.g., barcodes 5 and 9; see FIG. 2C,D) show a reduction in their relative proportions after BRAFi treatment (FIG. 2A, *stars*). Conversely, lineages in which many cells (>20%) reside in the fast-dividing cluster (e.g., barcodes 2 and 13; see FIG. 2C,D) significantly increase their relative proportions following treatment (FIG. 2A, *circles*). Indeed, a strong positive correlation exists (Pearson, R=0.55) between the proportion of cells in a lineage residing in the fast-dividing idling DTP state and the relative fold change of barcode abundance after treatment (FIG. 2E). In all, these results indicate that most clonal lineages survive BRAFi treatment and persist in the idling DTP state in similar proportions to the untreated population. Thus, rather than clonal selection, the experimental evidence points to all BRAF-mutant melanoma cells having the capacity to enter the idling DTP state, which is consistent with numerous recent reports on the behaviors and characteristics of DTPs (11–16).

**Figure 2:**
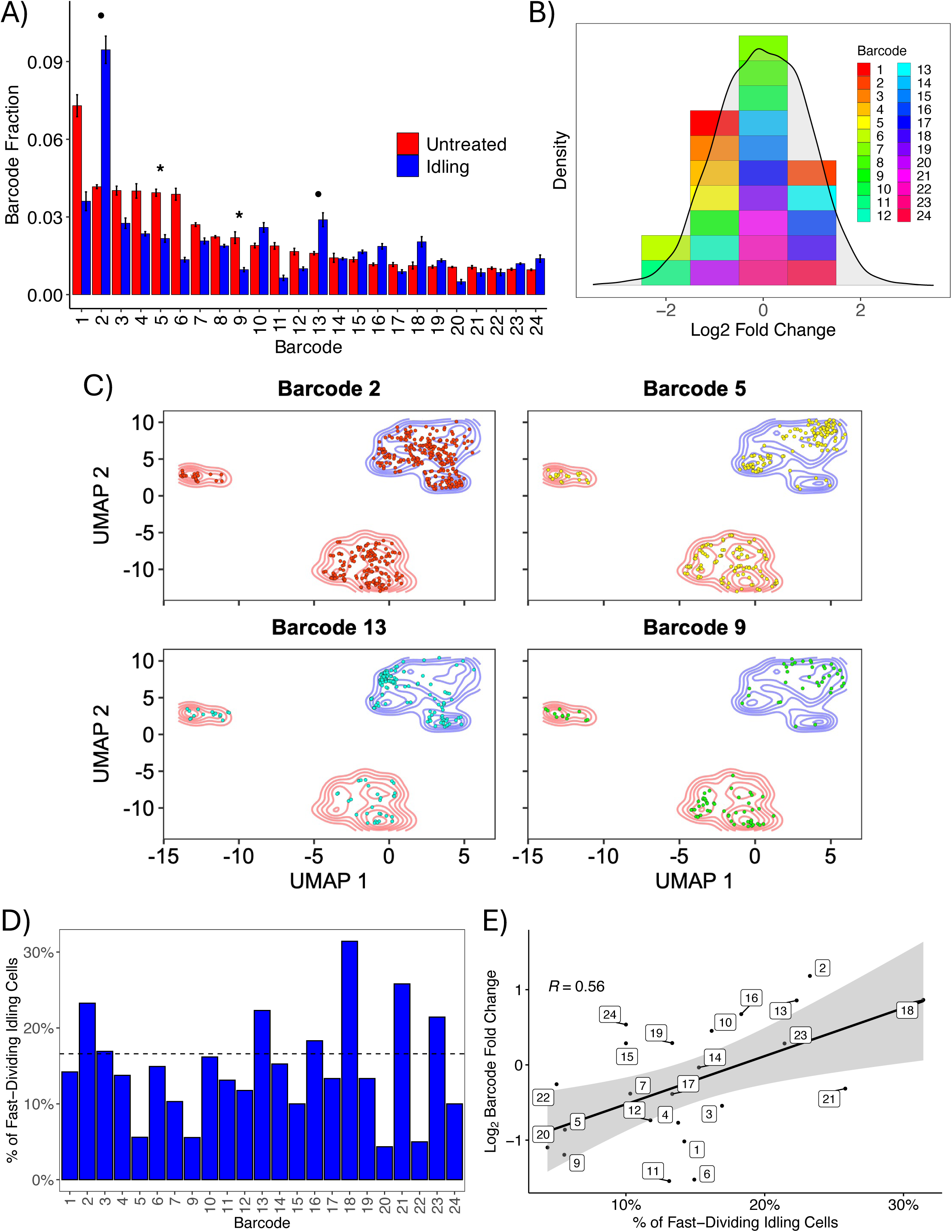
Most clonal lineages survive BRAFi treatment and enter the idling DTP state. **(A)** Relative fractions of the top 24 most abundant barcoded cell lineages in the untreated and idling DTP populations. Bar heights correspond to the average of three experimental replicates; lines are the standard deviations. Bars are ordered by abundance in the untreated condition. Stars (circles) signify example barcodes (plotted in *C*) with reduced (increased) abundance after treatment. **(B)** Distribution of (log2) fold-change from untreated to idling DTP state for barcoded clonal lineages. Means of fold-changes were compiled into a distribution for all captured lineages (grey) and the top 24 most abundant lineages from *A*. **(C)** Overlay of single cells from four barcoded lineages (stars and circles from *A*) on the transcriptomic UMAP space from FIG. 1A. Colored contours (red and blue) reflect cell density for both treatment conditions. **(D)** Proportion of cells in the fast-dividing (small) idling DTP cluster (see FIG. 1D) for the top 24 most abundant barcodes. The dashed line indicates the average for all barcodes. **(E)** Pearson correlation analysis between the percentage of cells in the fast-dividing idling cluster and the log_2_ fold-change in barcode abundance following BRAFi treatment for the top 24 most abundant barcodes.

### Multi-omics and calcium-flux analyses indicate changes in ion channel homeostasis and activity in drug-tolerant cells

We further analyzed the bulk RNA sequencing data from above for the three single-cell-derived BRAF-mutant melanoma subclones (FIG. 1B) by performing a differential expression analysis between the untreated (day 0) and idling DTP (day 8) populations. Inputting differentially expressed genes (DEGs) for all three subclones into a GO over-enrichment analysis returned numerous terms associated with ion transport and homeostasis (FIG. 3A). This led us to look at expression levels for numerous genes related to calcium handling (Materials and Methods). We see that many are upregulated or downregulated across the subclones after 3 and 8 days of BRAFi (FIG. 3B). These include genes encoding for sarcoendoplasmic reticulum Ca^2+^-ATPases (SERCAs; e.g., ATP2A1), inositol 1,4,5-trisphosphate receptors (ITPRs), stromal interaction molecule 1 (STIM1), and solute carrier (SLC) genes that encode for various membrane transport proteins. Additionally, we probed the epigenomic status of the idling DTP state by performing bulk ATAC (assay for transposase-accessible chromatin) sequencing (49) on samples of the same BRAF-mutant melanoma cell line, before and after 8 days of BRAFi treatment (FIG. S3A,B; see Materials and Methods). Regions of open chromatin on the DNA, referred to as “peaks,” were identified for both conditions (FIG. S3C). The distribution of binding loci (FIG. S3D) and corresponding peak feature set (FIG. S3E) show that idling peaks have fewer proximal features (e.g., promoter regions) and more distal elements (e.g., distal intergenic and intronic regions) compared to untreated cells. Distal elements have been known to be involved in short-term epigenetic regulation (50). Peaks unique to idling DTPs were then assigned to corresponding genes and these genes input into a GO over-enrichment analysis. As with the bulk RNA-sequencing data, numerous GO terms were returned associated with ion transport and activity (FIG. 3C). Indeed, GO terms from bulk RNA and ATAC sequencing data strongly correlated with each other, with the most significant terms being associated with ion-channel activity (FIGS. 3D and S3F,G). Thus, two independent data modalities (transcriptomics and epigenomics) provide strong evidence that alterations in ion-channel homeostasis and activity are characteristic of the idling DTP state.

To assess the functional consequences of these findings, we quantified calcium-channel activity in untreated and idling DTP cells using a calcium-flux assay that measures the amount of endoplasmic reticulum (ER)-resident calcium and the propensity for store-operated calcium entry (SOCE). In this assay, free ER calcium is measured first by using cyclopiazonic acid (CPA) in a calcium-free buffer to inhibit the activity of SERCAs, leading to the release of free calcium from the ER to the cytoplasm, where it is detected by a calcium-binding dye (FIG. 3E, *first peak*). Under normal conditions, cells undergo SOCE to replenish ER calcium upon depletion, i.e., ER calcium release stimulates the opening of plasma membrane-resident calcium channels (e.g., ORAI by STIM1), allowing extracellular calcium to flow into the cytoplasm before being pumped into the ER by SERCAs. Thus, to assess SOCE activity, calcium is added to the assay buffer after CPA addition and the subsequent increase in cytoplasmic calcium is quantified (FIG. 3E, *second peak*). Our results show that while untreated and idling DTP cells have essentially equal ER calcium content, SOCE activity in idling DTP cells is significantly reduced compared to untreated cells (FIG. 3E). These results verify that ion-channel activity is significantly altered in the idling DTP state.

**Figure 3:**
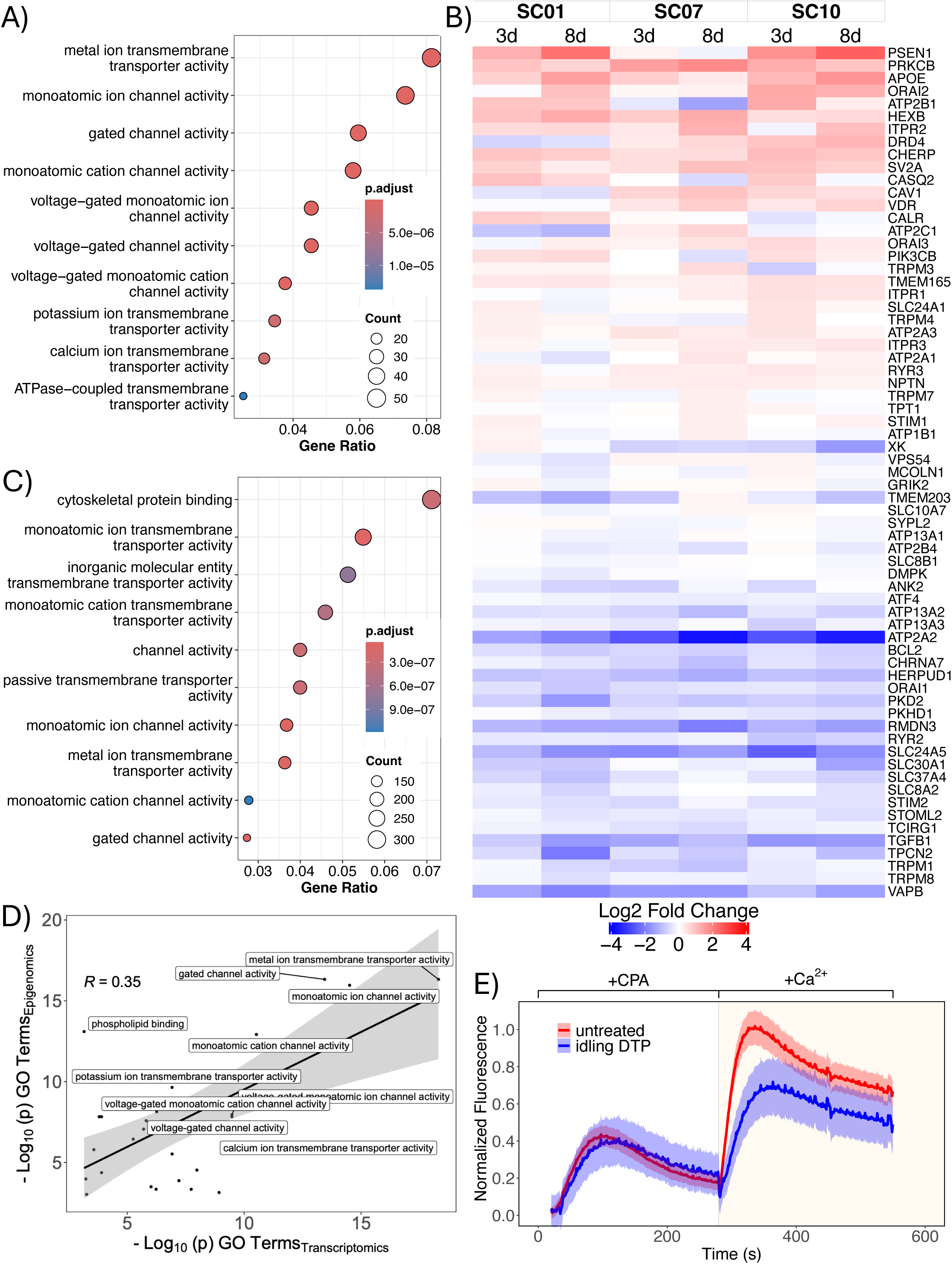
Idling DTPs are characterized by altered ion channel homeostasis and reduced calcium flux. **(A)** Gene ontology (GO) over-representation analysis of differentially expressed genes from bulk RNA sequencing of untreated and idling DTP populations. Shown are GO terms with the top 10 largest gene ratios. **(B)** Differential gene expression of calcium handling genes 3 and 8 days after BRAFi treatment in three single-cell-derived BRAF-mutant melanoma subclones. **(C)** GO over-enrichment analysis of genes associated with unique epigenomic peaks in idling DTP populations from ATAC sequencing. Shown are the top 10 terms with the largest gene ratio. **(D)** Correlation analysis between Molecular Function (MF)-type GO terms from transcriptomic data (panel *A*) and epigenomic data (panel *C*). **(E)** Calcium flux assay of untreated and idling DTP cells to assess endoplasmic reticulum calcium content/release and store-operated calcium entry (SOCE) activity. CPA: cyclopiazonic acid. Note that calcium is added to the culture medium in the presence of CPA and ionomycin is used to control for the number of cells in the assay (see Materials and Methods).

### Idling DTPs exhibit increased susceptibility to ferroptotic cell death

Previous reports indicate a complex interplay between ion channels, ER stress, and ferroptosis, a form of regulated cell death characterized by the iron-dependent accumulation of lipid peroxides, leading to oxidative damage to cell membranes (51–54). ER stress occurs when there is an imbalance between the protein-folding load in the ER and the ER’s folding capacity, which can be caused by various factors, including disturbances in calcium homeostasis (55). ER stress may, in turn, lead to increased cellular iron levels, which can promote the generation of ROS and lipid peroxidation, contributing to ferroptosis (29, 30). In our bulk RNA-sequencing data, we see altered expression of several genes related to ferroptosis in idling DTP cells (FIGS. 4A and S4). Notably, glutathione-metabolism gene expression is increased and genes for polyunsaturated fatty-acid (PUFA) enzymes are decreased in idling DTP cells. Interestingly, many DEGs feed into the Fenton reaction, which produces the free-radical precursors to ROS, the step directly before commitment to ferroptosis (FIG. S4).

To directly test susceptibility to ferroptosis of BRAF-mutant melanoma cell populations, we subjected both untreated and idling DTP cells to treatment with RSL3 (Materials and Methods), a compound that induces ferroptosis by targeting GPX4, a key regulator of glutathione oxidation and PUFA reduction (11). Interestingly, this treatment reduces glutathione metabolism while preventing the reduction of PUFA intermediates, leading to more precursors of the Fenton reaction and increased ROS, driving cell death by ferroptosis. We see that idling DTPs are ∼3-fold more sensitive to RSL3 by potency (IC50) than untreated cells (FIG. 4B). Furthermore, the addition of ferrostatin-1, a drug that inhibits the production of ROS by the Fenton reaction, rescues the enhanced sensitivity of the idling DTP population to RSL3 (FIG. 4C). These results indicate, therefore, that idling DTP cells are susceptible to death by ferroptosis induction, presumably because of increased ROS levels resulting from BRAFi-induced alterations to ion channel homeostasis and activity (FIG. 3). This is a potential vulnerability of idling DTPs that may be able to be exploited therapeutically.

**Figure 4:**
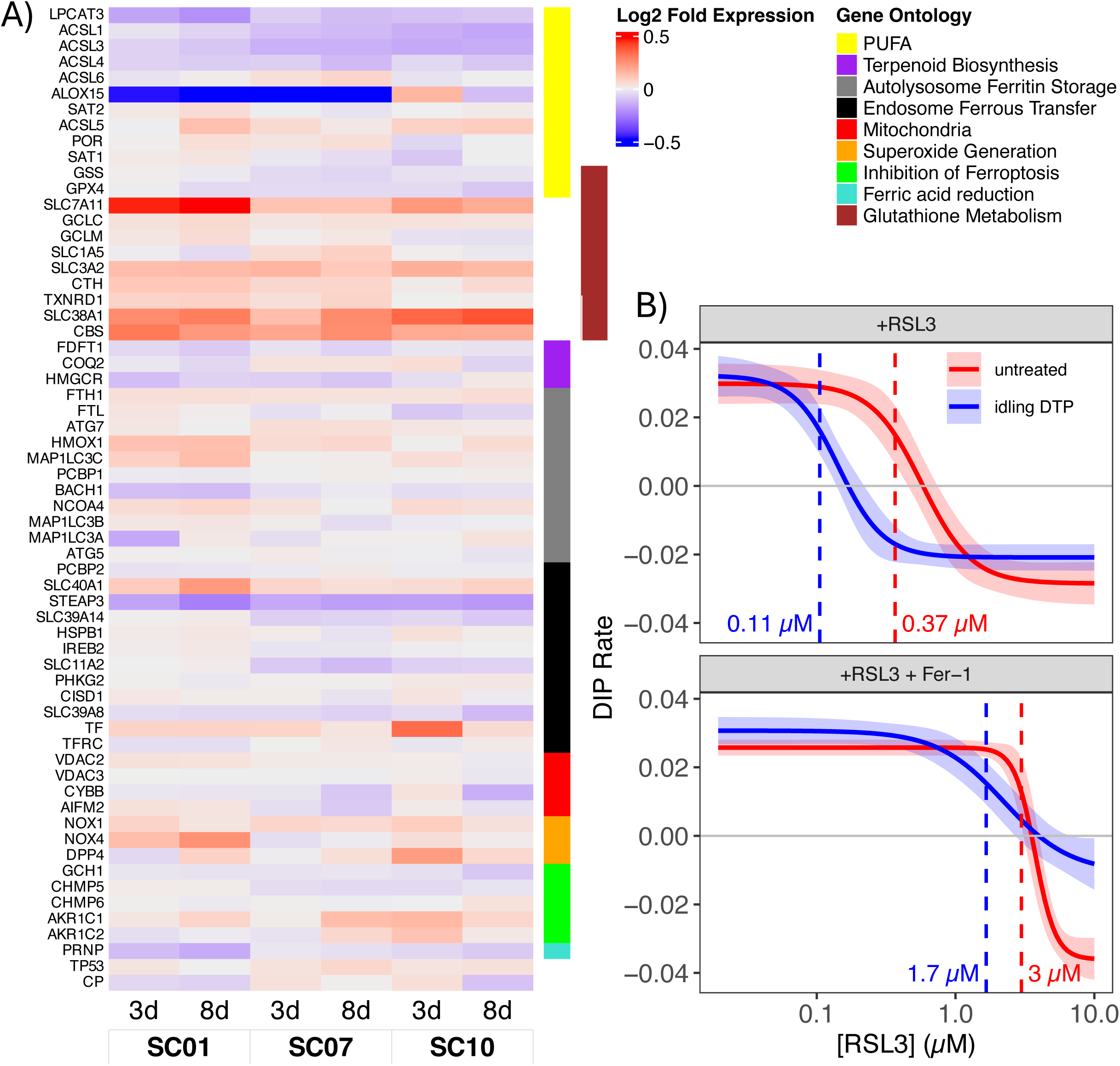
Idling DTP populations are susceptible to death by ferroptosis induction. **(A)** Gene expression values (log_2_ fold-change) at 3 and 8 days post-BRAFi treatment for three single-cell-derived subclones (SC01, SC07, SC10). Genes are grouped by ferroptosis-associated process (see FIG. S4). **(B)** Dose-response curves for untreated and idling DTP populations treated with the ferroptosis inducer RSL3 (*top*) and RSL3 together with ferroptosis inhibitor ferrostatin-1 (Fer-1; *bottom*). Dashed lines correspond to the half-maximal inhibitory concentration (IC_50_) for each condition, included in units of µM. DIP: drug-induced proliferation (see Materials and Methods).

## DISCUSSION

Drug tolerance has been observed and described in many cancer types (14, 16, 27, 37, 56–58), yet the processes that give rise to DTPs have been interpreted in various ways (57). Some studies report treatment-induced protective programs in which drug exposure rapidly triggers new transcriptional or epigenetic responses that were not evident before treatment (12, 14, 16, 59). Others emphasize rare, pre-existing cell types that differ from the bulk population and become enriched under therapy (13, 15, 60). A third group of studies highlights stochastic, reversible phenotypic states that cells already visit before treatment but become stabilized once drug is applied (6, 14, 16, 60–62). These varying perspectives emphasize different origins for drug tolerance but acknowledge that treatment exposure changes which phenotypic states persist within a cancer cell population and that multiple drug-tolerant states may co-exist (27, 37, 56, 58, 63–65). However, these prior reports also describe DTPs, at least implicitly, as uniform states in which each cell within the population is essentially identical.

In this work, we build on these prior studies but provide evidence for a fundamentally different view of the nature of DTPs, i.e., not as uniform phenotypes but rather as heterogeneous collections of phenotypic sub-states that, together, achieve near-zero net population growth under treatment conditions. Specifically, across multiple assays, untreated and idling DTP populations in BRAF-mutant melanoma exhibited significant population-level differences, consistent with prior reports (17, 20). In bulk and single-cell transcriptomic data, the idling DTP population showed reduced but still appreciable heterogeneity relative to the untreated population (FIG. 1). This treatment-exposed population occupied a narrower range of phenotypic states, some of which showed distinct functional programs and some that partially overlapped with those observed in untreated conditions (FIG. 1C). Lineage tracing indicated that drug tolerance does not arise from a small set of pre-existing lineages but rather emerges from cells in virtually all lineages (FIG. 2). Single-cell RNA-seq, bulk RNA-seq, and bulk ATAC-seq, together with GO analyses, identified enrichment of ion-channel and calcium-handling pathways, consistent with treatment-associated changes observed in calcium-flux assays (FIG. 3) and with similar treatment-associated shifts in calcium regulation reported in previous melanoma studies (18, 35, 36, 55, 66). Consistent with these transcriptional and functional changes, the idling DTP population exhibited increased sensitivity to ferroptosis induction relative to the untreated population (FIG. 4). Similar vulnerabilities have been reported in other cancer systems, including ferroptosis sensitivity of DTPs and therapy-modulated redox and lipid metabolism (11, 29, 30, 52–54).

It is important to note that a limitation of the present study is that our data represent population snapshots rather than time-lapse trajectories. Thus, we do not directly observe how individual cells move through the landscape over time. Additionally, our ferroptosis experiments establish a population-level vulnerability in this melanoma model but do not identify the specific molecular drivers of that susceptibility, and we cannot assume all DTP populations will share the same vulnerability (11, 30, 52–54). We also did not isolate and analyze drug-tolerant populations following ferroptosis induction, which are also expected to be composed of multiple phenotypic sub-states. Nonetheless, the observations reported here build on prior work (17) and provide substantial experimental support for the view that drug treatment modifies the epigenetic landscape on which cancer cells reside. The population re-equilibrates across this drug-modified landscape that, while less heterogeneous than the untreated landscape, still comprises multiple phenotypic states (FIG. 5A). Moreover, cells from all parts of the untreated landscape transition into the DTP sub-states, which are characterized by distinct molecular processes (20), such as increased oxidative phosphorylation and decreased calcium flux (FIG. 5A). This perspective helps unify diverse observations in the DTP literature (57, 67–70). For example, treatment-induced programs, persistence of pre-existing lineages, and stabilization of fluctuating phenotypes (13–16, 60) can all be viewed as different ways in which a population adjusts to treatment-driven phenotypic changes, rather than fundamentally distinct mechanisms of drug tolerance emergence. The co-existence of multiple DTPs within a drug-tolerant tumor can also be accommodated within this conceptual framework. Not only can multiple DTP states co-exist, across which cells can transition, but each is itself a collection of traversable sub-states that either expand or deplete the total cell population (FIG. 5B).

This view adds an additional layer of complexity to the issue of DTP heterogeneity, with consequences for future therapies aimed at eradicating drug-tolerant populations. The current philosophy for targeting DTPs focuses on the isolation and molecular characterization of DTP states to identify actionable vulnerabilities, under the assumption that the DTP population is phenotypically homogeneous (21–28). Within the framework proposed here, this approach will be unsuccessful because the phenotypic landscape that cancer cells occupy is heterogeneous and adjusts in response to perturbations. This provides a means of escape, whereby cells transition into remaining or emergent viable sub-states. A game of “whack-a-mole” provides a useful metaphor for this scenario, where a player constantly chases the mole but is unable to strike it because it moves to a new location with each whack. Rather than engaging in this futile pursuit, a more fruitful strategy would be to change the rules of the game and reshape the board itself, e.g., by filling holes or altering its slope, thereby blocking avenues for the mole’s escape. Analogously, a more durable approach to treating DTPs would be to alter the phenotypic landscape on which the cells reside. This “targeted landscaping” strategy would entail multiple sequential interventions to progressively modify the topography of the landscape to favor regressing over expanding sub-states, until the entire DTP population is eliminated (FIG. 5C). By shifting the focus to how treatment reshapes the phenotypic landscape and the nature of the sub-states within it, the framework proposed in this work offers a generalizable path toward controlling, and potentially eliminating, DTPs before acquired treatment resistance arises.

**Figure 5.**
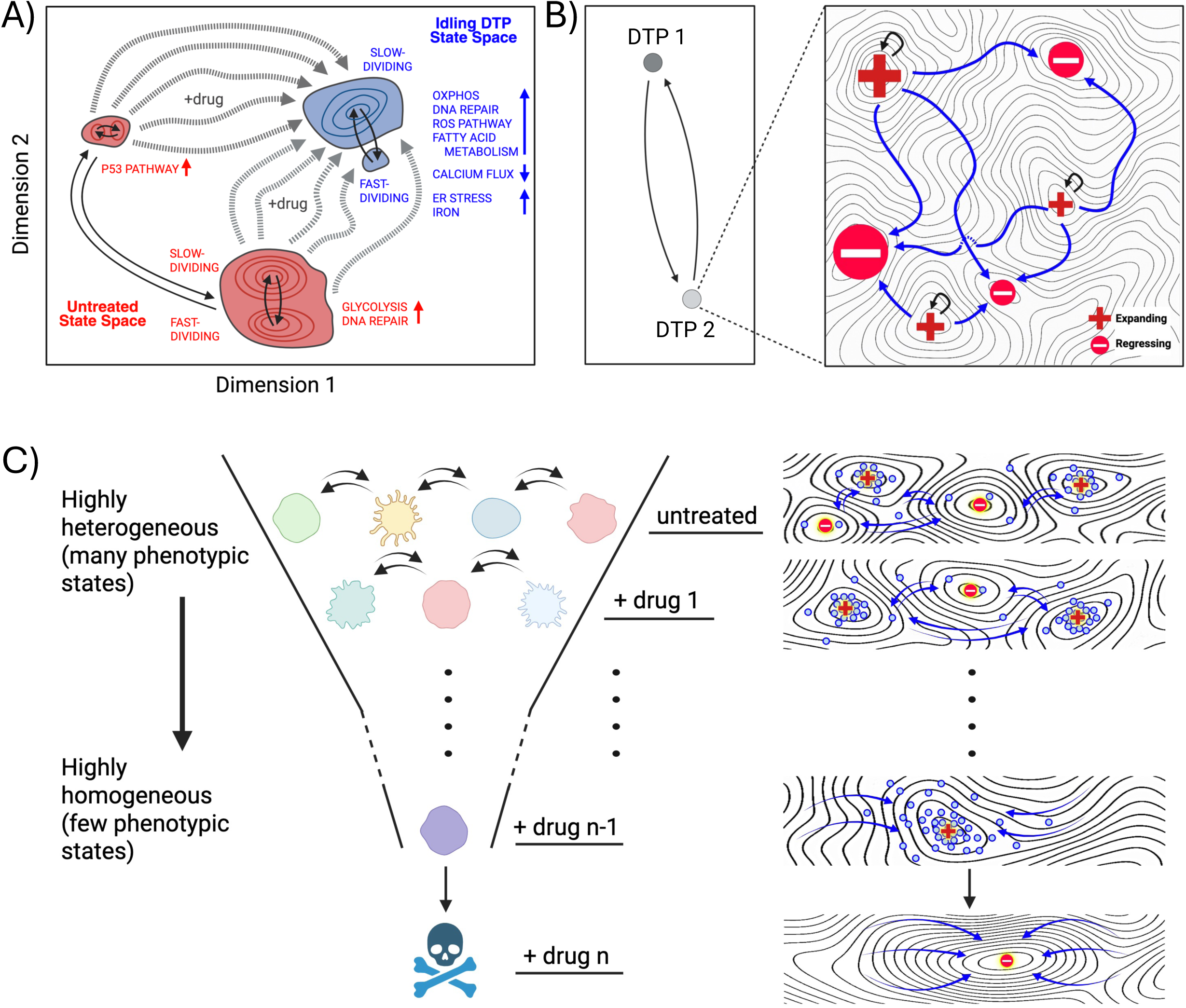
Conceptual framework describing drug-induced phenotypic landscape reshaping, DTP heterogeneity, and targeted treatment strategies. (**A**) Illustrative interpretation of the drug-induced transition of a BRAF-mutant melanoma population to the idling DTP state. The clusters in the UMAP space correspond to those observed experimentally (FIG. 1A). Within the framework proposed in this work, the untreated phenotypic landscape (*red*) comprises multiple, disparate sub-states across which cells can transition (*solid arrows*). BRAFi treatment modifies the landscape, reducing but not eliminating phenotypic heterogeneity (*blue*). Cells from any untreated sub-state can transition into the idling DTP population state (*dashed arrows*) and occupy any of its sub-states, across which cells can transition (*solid arrows*). Consistent with GO analyses (FIGS. 3A,C and S1B,C), different phenotypic sub-states within the untreated and idling DTP populations exhibit distinct functional programs, including metabolic pathways, ion channel activity, and stress response processes. **(B)** Illustration of multiple levels of DTP heterogeneity. (*Left*) Consistent with prior reports, multiple distinct DTP states can coexist within a tumor (“DTP 1” and “DTP 2”), among which cells can transition. (*Right*) Novel to the framework presented here, each DTP state is a collection of phenotypic sub-states, with proliferation rates ranging from positive (“Expanding”) to negative (“Regressing”). Cells dynamically transition across these sub-states until a steady-state is reached, in which the population as a whole exhibits a near-zero net proliferation rate. **(C)** Hypothetical “targeted landscaping” treatment strategy. Based on the framework presented in this work, sequential therapies are applied to reshape the phenotypic landscape and reduce the available phenotypic sub-states within the DTP population state. Ultimately, the goal is to produce a phenotypic landscape composed entirely of regressing states, resulting in the elimination of all drug-tolerant cells. All panels were created with BioRender.com.

## MATERIALS AND METHODS

### Cell culture and reagents

The SKMEL5 cell line, chosen for this study because it exhibits median BRAFi sensitivity compared to other BRAF-mutant melanoma cell lines (36), was purchased from the American Type Culture Collection (ATCC) and labeled with a fluorescent histone H2B conjugated to either the monomeric red fluorescent protein (H2BmRFP) or the green fluorescent protein (H2B-GFP), as previously described (6, 17). Cells were initially stored at -80°C and then moved into liquid nitrogen. Single cell-derived subclones of SKMEL5 were selected and derived by limiting dilution, as described previously (17). Cells were cultured in a mixed media (1:1) of Dulbecco’s Modified Eagle Medium (DMEM) and Ham’s F-12 medium (both from Gibco, Grand Island, NY, USA), supplemented with 10% fetal bovine serum (FBS) (Thermo Fisher Scientific, Waltham, MA, USA), incubated at 37°C in 5% CO_2_, and passaged twice a week using TrypLE (Gibco, Grand Island, NY, USA). Cells were also tested for mycoplasma contamination using the MycoAlert^TM^ mycoplasma detection kit (Lonza, Basel, Switzerland), according to the manufacturer’s instructions, and confirmed to be mycoplasma-free. SKMEL5-H2BmRFP cells were further transduced with a cellular barcoding library (see “Cellular barcoding”) using HEK293T cells (Thermo Fisher Scientific, Waltham, MA, USA) to produce lentiviral vectors.

SKMEL5 cells were treated with either dimethyl sulfoxide (DMSO; vehicle control) or, to induce them into the idling DTP state, 8 μM of the BRAFi PLX4720 (research analog to Vemurafenib) for eight days, with media/drug changed every three days. These cells were used for all subsequent experiments and analyses. PLX4720, ferroptosis inducer RSL3, and ferroptosis inhibitor ferrostatin-1 (Fer-1) were obtained from MedChem Express (Monmouth Junction, NJ, USA) and solubilized in DMSO at a stock concentration of 10mM and stored at -20°C. Bovine serum albumin (BSA; Research Products International, Mt. Prospect, IL, USA) and phosphate-buffered saline (PBS; Corning Inc., Corning, NY, USA) were used to prepare cells for single-cell RNA sequencing. Detergents NP-40, Tween20, and Digitonin (MilliporeSigma, Burlington, MA, USA) were used to lyse cells and permeabilize nuclei in preparation for bulk ATAC sequencing. For calcium flux assays, we used calcium chloride (CaCl_2_; Thermo Fisher Scientific, Waltham, MA, USA, Cat #: J63122.AE) as the source of Ca^2+^, the Ca^2+^ ionophore ionomycin (Cayman Chemical Company, Ann Arbor, MI, USA, Cat #: 11932), green-fluorescent calcium binding dye Fluo-8-AM (AAT Bioquest, Pleasanton, CA, USA), and Hanks’ Balanced Salt Solution (HBSS; no Ca^2+^; Gibco, Grand Island, NY, USA, Cat #: 14175-095) supplemented with 10 mM HEPES buffer (Corning Inc., Corning, NY, USA, Cat #: 25-060-CI).

### Cellular barcoding

#### Setup

The cellular barcoding library was constructed by cloning a guide RNA (gRNA) library of barcodes into a CROP-seq-BFP-TSO vector, as previously described (43, 44). The vector was engineered such that barcodes can be found by isolation and amplification (barcode sampling) or mRNA capture in a single-cell RNA sequencing (scRNA-seq) experiment. gRNAs were built as a 20-nucleotide sequence of four nucleotides identical among all barcodes (XXXX), followed by 16 strong-weak (SW) paired nucleotides (i.e., XXXXSWSWSWSWSWSWSWSW). The SW pairing of the barcode sequence was designed to prevent polymerase chain reaction (PCR) amplification bias. The maximum complexity of this library is 2^16^ (∼65,000 unique barcodes). The barcode library vector was used to produce lentiviral libraries using a lipofectamine transfection of HEK293T cells. Media containing lentiviral particles were collected at 48- and 72-hours post-transfection, pooled, filtered through a 0.45 µm Nalgene syringe filter (Thermo Fisher Scientific, Waltham, MA, USA), and concentrated using a 50 mL size-exclusion column (MilliporeSigma, Burlington, MA, USA) by centrifugation at 2200 relative centrifugal field (RCF) at 4°C for two hours. Concentrated virus was stored in -80°C. SKMEL5 cells were seeded in a 6-well plate at ∼1x10^6^ cells per well in 2.5 mL culture media. Cells were transduced with the barcoded CROP-seq-BFP-TSO-Barcode_sgRNA lentivirus using 0.8 mg/mL in each well and a multiplicity of infection (MOI) of 0.05, as previously described (43, 44). Twenty-four hours after incubation, transduction media (containing polybrene) was exchanged for fresh culture media. Forty-eight hours after incubation, barcoded cells were isolated by fluorescence-activated cell sorting (FACS) and subsequently cultured until confluence in a T-150 dish and cryopreserved. Cells were then thawed in a T-25 dish and scaled up for ∼2 weeks in two separate sets. The first set of thawed cells was treated with 8 µM PLX4720 for eight days and subjected to barcode sampling (see “Cellular Barcoding: *Barcode Sampling Analysis*”), along with an untreated control sample. The second set of thawed cells was plated in three T-75 flasks (parallel replicates) and independently treated with 8 µM PLX4720 for eight days and subjected to scRNA-seq (along with an untreated control sample) by the 10X genomics Chromium platform (version 2 chemistry; see “Single-cell RNA transcriptomic sequencing”). For all drug-treated samples, media and drug were replaced every three days. Untreated cells were expanded completely over the entire time course (no cell splitting).

#### Barcode Sampling

For the first set of thawed cells (see “Cellular Barcoding: *Setup*”), untreated and 8-day PLX4720-treated cells were pelleted for gDNA extraction using the DNeasy Blood and Tissue Kit (Qiagen, Hilden, Germany), per manufacturers’ instructions. Barcode sequences were amplified for each replicate by PCR (98°C for 30 seconds, followed by 22 cycles of denaturation at 98°C for 10 seconds, annealing at 63°C for 30 seconds, extension at 72°C for 10 seconds, and a final extension at 72°C for 5 minutes) using primers containing flanking regions and Illumina adapter index sequences (Illumina, San Diego, CA, USA). For each PCR reaction, 2 µg gDNA was used and a combination of five distinct pooled forward primers was utilized to minimize sequencing error. Reactions were purified using a 1.8x AMPure XP bead (Beckman Coulter, Brea, CA, USA) cleanup. Reaction products were confirmed using agarose gel confirmation (band at ∼215 bp). The resulting libraries were quantified using a Qubit fluorometer (Thermo Fisher Scientific, Waltham, MA, USA), Bioanalyzer 2100 (Agilent Technologies, Santa Clara, CA, USA) for library profile assessment, and quantitative PCR (qPCR; Kapa Biosystems, Wilmington, MA, USA, Cat #: KK4622) to validate ligated material, according to the manufacturer’s instructions. The libraries were sequenced using the NovaSeq 6000 with 150 bp paired-end reads as sequencing spike-ins (targeting ∼200,000 reads). ‘Real-Time Analysis’ (‘RTA’) software (version 2.4.11; Illumina, San Diego, CA, USA) was used for base calling and ‘MultiQC’ (version 1.7) (71) was used for quality control. Barcodes were identified from amplified sequence reads by trimming flanking adapter sequences (5’: XXXX; 3’: XXXX). Barcode abundances were totaled and normalized to the library read depth, resulting in reads per million (RPM). Barcodes less than 100 counts per million (CPM) were removed from the analysis. Numbers of unique barcodes were calculated based on this threshold.

### Single-cell RNA transcriptomic sequencing

#### Data Collection

For the second set of thawed cells (see “Cellular Barcoding: *Setup*”), untreated and 8-day PLX4720-treated cells were prepared targeting ∼3000 cells per sample, washed, and resuspended in 0.04% BSA in PBS. Cell suspensions were subjected to a 10X Genomics single-cell gene expression protocol (version 2, 3’ counting; 10xgenomics.com/support/single-cell-gene-expression/documentation/steps/ library-prep/assay-scheme-and-configuration-of-chromium-single-cell-3-v-2-libraries) in two separate wells, according to the manufacturer’s guidelines. Single-cell mRNA expression libraries were prepared according to the manufacturer’s instructions. Due to the nature of the gRNA barcoding library construction (43, 44), mRNAs resulting from gRNA barcodes were captured along with other mRNAs. Libraries were cleaned using solid-phase reversible immobilization (SPRI) beads (Beckman Coulter, Brea, CA, USA) and quantified using a Bioanalyzer 2100 (Agilent Technologies, Santa Clara, CA, USA). The libraries were sequenced using the NovaSeq 6000 (Illumina, San Diego, CA, USA) with 150 bp paired-end reads targeting 50 million reads per sample for the mRNA library (including the barcode library). ‘RTA’ software (version 2.4.11; Illumina, San Diego, CA, USA) was used for base calling and ‘MultiQC’ (version 1.7) (71) for quality control. Gene counting, including alignment, filtering, barcode counting, and unique molecular identifier (UMI) counting was performed on each library using the ‘*count’* function in the 10X Genomics software ‘Cell Ranger’ (version 3.0.2) (72) with the GRCh38 (hg38) reference transcriptome (73).

#### Transcriptome Analysis

We used ‘sc-UniFrac’ (61) to quantify the degree of overlap between the two scRNA-seq count matrices (untreated and idling DTP) obtained from ‘Cell Ranger’ (see “Single-cell RNA transcriptomic sequencing: *Data Collection*”). Since overlaps were minimal and cells were prepared and processed in parallel, no computational batch correction was performed. We used ‘Seurat’ (version 3) (74) to perform gene expression analysis. The ‘*SCTransform’* function was used to regress out mitochondrial gene expression (‘percent.mt’), number of features (i.e., genes; ‘nFeature_RNA’), number of RNA molecules in the cell (‘nCount_RNA’), and cell cycle variables (‘S.Score’ and ‘G2M.Score’). Feature selection was performed according to ‘Seurat’ guidelines using a variance stabilizing transformation of the top 2000 most variable features. Data was normalized and scaled according to ‘Seurat’ guidelines. Data between conditions were combined and visualized using the UMAP dimensionality reduction algorithm (31), as implemented in ‘Seurat’. Clustering was performed in the joint UMAP space using the default ‘Seurat’ implementation, a shared nearest neighbor (SNN) modularity optimization-based method (75). Differential expression analysis was performed using the ‘*FindMarkers’* function in ‘Seurat’. Differentially expressed genes (DEGs; adjusted-p < 0.05) were input into a GO over-enrichment analysis (38).

#### Barcode Analysis

After calculating the scRNA-seq gene expression matrices with ‘Cell Ranger’ (see above), barcode abundances were incorporated into the matrices. To do this, gRNA lineage barcodes were mapped to their associated 10X cell barcodes. First, unmapped scRNA-seq Binary Alignment Map (BAM) files were cleaned to only include the mRNA transcript ID, scRNA-seq cell barcode, and scRNA-seq UMI. Mapped scRNA-seq BAM files (3’ heavy) were cleaned to only include the mRNA transcript ID and lineage barcode (from the gRNA library). Unmapped and mapped subsets were then merged on the mRNA transcript ID to assign a lineage barcode to each cell barcode and UMI. The resulting merged dataset was paired down to a cell barcode/lineage barcode pair, which was appended to each cell in the gene expression matrix as a metadata tag. Barcode abundances were totaled across all cells in the experiment that captured a barcode. Barcodes overlaid on UMAP projections of scRNA-seq data were further categorized into “fast-dividing” (G1) and “slow-dividing” (S-G2-M) states based on cell cycle score (39) (see “Single-cell RNA transcriptomic sequencing: *Transcriptome Analysis*”).

#### Functional Interpretation Analysis

The single-cell transcriptome count matrices obtained from ‘Cell Ranger’ (see “Single-cell RNA transcriptomic sequencing: *Data Collection*”) were scaled by multiplying counts by the median RNA molecules across all cells and dividing that number by the number of RNA molecules in each cell. Hallmark gene sets (50 in total) were downloaded from the molecular signatures database (MSigDB) (32–34). The scaled count matrix and each hallmark gene set were input into ‘VISION’ (76) to identify gene signature scores for each cell-signature pair. Four hallmark gene sets (KRAS_SIGNALING_UP, KRAS_SIGNALING_DOWN, UV_RESPONSE_UP, UV_RESPONSE_DOWN) were condensed into two (KRAS_SIGNALING, UV_RESPONSE) by ‘VISION’ to leave 48 total gene signatures. Scores were compiled into a distribution and plotted by cluster (k=3) for each gene set.

### Bulk RNA transcriptomic sequencing

#### Data Acquisition

Total RNA was isolated from untreated and BRAFi-treated SKMEL5 single cell-derived subclones, each in triplicate, using the TRIzol isolation method (Invitrogen, Carlsbad, CA, USA), according to the manufacturer’s instructions. RNA samples were submitted to Vanderbilt VANTAGE Core services for quality check, where mRNA enrichment and cDNA library preparation were done with Illumina TruSeq stranded mRNA sample prep kit (Illumina, San Diego, CA, USA). Sequencing was done at Paired-End 75 bp on the Illumina HiSeq 3000. Reads were aligned to the hg38 human reference genome (73) using ‘HISAT2’ (77) and gene counts were obtained using ‘featureCounts’ (78).

#### Data Analysis

RNA-seq data was analyzed using the ‘DESeq2’ R package (79). Cells with less than 18 reads per condition were removed, according to ‘DESeq2’ vignettes. Counts were transformed using the regularized logarithm (‘*rlog’*) normalization algorithm. PCA was performed using the ‘*prcomp’* function in R (rdrr.io/r/stats/prcomp.html). Differential expression analysis was performed using a ‘DESeq2’ model design to quantify both changing variables and their interaction (∼ subline + treatment time + subclone:treatment time). DEGs across subclones between untreated (pre-treatment, day 0) and idling DTP (post-treatment, day 8) were identified (adjusted-p < 0.05, log2 fold-change > 2) and input into a GO enrichment analysis (38), which identified GO terms associated with “Biological Process” (BP), “Molecular Function” (MF), and “Cellular Component” (CC) GO categories.

#### Calcium Handling Gene Signature Analysis

Using the ‘AmiGO’ web application (80) genes associated with calcium handling were identified by the GO term “intracellular calcium ion homeostasis” (Accession #: GO:0006874; Release date: 2023-11-17; Version #: 10.5281/zenodo.10162580). Inositol 1,4,5 trisphosphate receptors (ITPRs) and other transient receptor potential melastatin (TRPM) receptors were also added manually to the gene set.

#### Ferroptosis Gene Signature Analysis

A ferroptosis gene signature was obtained from the WikiPathways database (https://wikipathways.org/pathways/WP4313.html) (81, 82). For each single cell-derived subclone, mRNA levels were normalized to the 0-day time point and a log_2_ fold-change was calculated compared to the 0-day baseline.

### Bulk ATAC epigenomic sequencing

#### Data Acquisition

Data was collected using the Omni-ATAC protocol (83) for bulk assay for transposase-accessible chromatin with sequencing (ATAC-seq). For the second set of thawed cells (see “Cellular Barcoding: *Setup*”), untreated and 8-day PLX4720-treated cells were pelleted at ∼50,000 cells and resuspended in a cold ATAC-seq resuspension and lysis buffer containing NP-40 (0.1%), Tween20 (0.1%), and Digitonin (0.01%) and incubated on ice. A resuspension buffer was added (0.1% Tween20, no NP-40 or Digitonin) to wash out the lysis reaction. Cells were pelleted and resuspended in a transposition mix (5x Tris-DMF, phosphate-buffered saline (PBS), 1% Digitonin, 10% Tween20, nuclease-free H_2_O), including transposase Tn5, followed by a 30-minute incubation at 37°C, with shaking to enhance tagmentation. After 30 minutes, the reaction was stopped by adding a DNA binding buffer (Zymo Research, Irvine, CA, USA) and purified using a DNA Clean and Concentrate kit (Zymo Research, Irvine, CA, USA, Cat #: D4004). The final product was eluted in nuclease-free H_2_O. PCR amplification was performed on the eluate with an NEBNext High-Fidelity 2X PCR Master Mix (New England Biolabs, Ipswich, MA, USA, Cat #: M0541S) and custom primer indexes (extension at 72°C for 5 minutes; denaturation at 90°C for 30 seconds; 12 cycles: denaturation at 98°C for 10 seconds, annealing at 62°C for 30 seconds, extension at 72°C for 30 seconds; final extension at 72°C for 5 minutes). The PCR product was purified with the Zymo DNA Clean and Concentrate kit and eluted in 22 mL nuclease-free H_2_O. ATAC-seq PCR libraries were visualized by agarose gel electrophoresis for a quick check for the nucleosome ladder pattern (bands over ∼150 bp). Libraries were also quantified using a Qubit fluorometer (Thermo Fisher Scientific, Waltham, MA, USA), Bioanalyzer 2100 (Agilent Technologies, Santa Clara, CA, USA) for library profile assessment, and qPCR (Kapa Biosystems, Wilmington, MA, USA, Cat #: KK4622) to validate ligated material, according to the manufacturer’s instructions. The libraries were sequenced using the NovaSeq 6000 with 150 bp paired-end reads as spike-ins on the sequencing chip (untreated: ∼160 million reads, idling DTP: ∼130 million reads). ‘RTA’ software (version 2.4.11; Illumina, San Diego, CA, USA) was used for base calling and ‘MultiQC’ (version 1.7) (71) for quality control.

#### Data Analysis

Reads were trimmed using ‘cutadapt’ (84) (paired-end) to remove primer sequences and aligned to the hg38 reference genome (73) using the ‘*bwa-mem’* function in ‘Burrows-Wheeler Aligner’ (BWA; version 0.7.17) (85). Aligned reads were sorted and duplicates were marked using the ‘Picard’ software tool (version 2.17.10; broadinstitute.github.io/picard). Untreated reads had more detected duplicates (∼78% compared to ∼32% in idling DTPs). Reads were deduplicated, but to address the discrepancy in deduplicated library complexity, idling DTP reads were subsampled (25% of the original library) to achieve a similar complexity to the untreated sample (FIG. S3A,B). Insert sizes were plotted from the output of ‘*InsertSizeMetrics’* in ‘Picard’ after deduplication. Peaks of open chromatin were called using the *‘callpeak’* function in ‘MACS2’ (86), according to recommended guidelines for ATAC-seq data (BAM paired-end method, q-threshold: 0.05, no ‘MACS2’ model, shift: -100, extension size: 200). Peaks were subjected to a further round of quality control and cleaning using ‘ChIPQC’ (87) (peak mapping, peak duplication, blacklist peak detection) and blacklisted peaks were removed. Peaks were converted to consensus counts using the ‘*runConsensusCounts’* function in ‘soGGi’ (rdrr.io/bioc/soGGi/). Intersections and unique cleaned peaks were determined and visualized as a Venn diagram (FIG. S3C) using the ‘*vennDiagram’* function in the ‘limma’ software package (88). Unique and intersection peaks were annotated with nearest neighbor genes using the ‘*annotatePeak’* function and the hg38 transcriptome (73) in the ‘ChIPseeker’ software package (89). These peaks were also re-aligned to the transcription start site (TSS) for each gene (FIG. S3D). Peaks were classified based on closeness to the TSS and assigned to a predicted feature (promoter, UTR, exon, intron, downstream, distal intergenic; FIG. S3E). Genes associated with unique peaks and intersections of peaks were also input into a GO enrichment analysis for BP, MF, and CC GO categories (same as “Bulk RNA Transcriptomic Sequencing: *Data Analysis*” above).

### Gene ontology analysis

Genes associated with unique ATAC-seq peaks (see “Bulk ATAC Epigenomic Sequencing: *Data Analysis*”) and DEGs from bulk RNA-seq (for all single cell-derived subclones; see “Bulk RNA Transcriptomic Sequencing: *Data Analysis*”) were identified for untreated and idling DTP cell populations. The two gene lists were independently subjected to a GO enrichment analysis using ‘clusterProfiler’ (90, 91) and genes were compared to BP, MF, and CC GO categories. For each category, GO terms significantly enriched in the unique ATAC-seq peaks (p < 0.05) and in DEGs (p < 0.05) were identified and stored independently as separate GO term lists. For all GO terms shared between the two lists, we calculated -log_10_(p-value) and ranked terms based on statistical significance. Spearman correlation (92) was calculated between the significant GO terms using ‘ggpubr’ (version 0.4.0; rpkgs.datanovia.com/ggpubr).

### Calcium flux assays

#### Data Acquisition

Barcoded SKMEL5 cells were plated onto a 384-well tissue-culture-treated plate 24 hours before imaging at a density of 10,000 cells/well. For the treated condition, cells were treated with 8 µM PLX4720 for eight days, with fresh media/drug swapped out every three days. Cells were trypsinized, counted, and seeded onto imaging plates, all while in the continuous presence of 8 µM PLX4720. Untreated control cells were taken from the same cell line culture, which was maintained separately. On the day of experimentation, the 384-well plate with all treated and untreated cells was incubated with 4 µM Fluo-8-AM in fresh culture media (10% FBS) for 1 hour at room temperature, as recommended by the manufacturer. Dye-containing medium was removed and HBSS (10 mM HEPES, no Ca^2+^) was used to wash the wells of excess dye, followed by removal and addition of 20 µL of fresh HBSS for use as the assay buffer. The plate was loaded into a Panoptic kinetic imaging plate reader (WaveFront Biosciences, Franklin, TN, USA), which combines liquid handling capabilities and the ability to record fluorescent intensity from each well of the 384-well plate. A custom three-addition protocol was developed to add the various drugs and assay conditions to the plate during the SOCE assay. Drug-addition reservoir plates were loaded with assay buffer (HBSS, with or without Ca^2+^) and thoroughly mixed immediately before imaging. Add conditions were divided into three parts: 1) cyclopiazonic acid (CPA; final concentration of 50 µM) to inhibit the activity of Sarcoendoplasmic Reticulum Calcium ATPase (SERCA), leading to ER Ca^2+^ release; 2) addition of Ca^2+^ to the assay condition to activate SOCE activity; and 3) addition of ionomycin, the Ca^2+^ ionophore (final concentration of 5 µM), to generate maximal signal intensity to control for variations in cell count in individual wells (this was particularly important since drug-treated idling DTP cells experienced increased washout due to the stressful nature of sustained BRAFi treatment). Fluo-8-AM was excited at 480 nm and imaged at 538 nm, with a frequency of 1 Hz. The CPA treatment condition was imaged for 260 seconds before the addition of Ca^2+^, followed by 270 seconds of imaging before the addition of 5 µM ionomycin. Treatment conditions were replicated in sets of eight and average values traced with 95% confidence intervals.

#### Data Analysis

For each condition (untreated and idling), fluorescence data were normalized by subtracting from each well the lowest average intensity value near the start of the experiment (t < 10 s) and then dividing by the highest average intensity value during the ionomycin treatment phase (t > 530 s) to control for the number of cells in each treatment group (the idling DTP population tended to have fewer cells than the untreated population). For comparing calcium flux between the treatment groups, means and standard errors (σ/√N) were calculated for each time point and plotted using ‘ggplot2’ (version 3.2.0) (93). For visualization purposes, the earliest time points (t < 10 s) and data from the ionomycin treatment phase (t > 530 s) were excluded from the plots.

### Ferroptosis-induction experiments

Plates of H2B-GFP-labeled SKMEL5 cells (see “Cell culture and reagents”) were treated with either vehicle (DMSO) or BRAFi (8 μM PLX4720) for eight days, incubated at 37°C and 5% CO_2_, changing media (with vehicle/drug) every three days. BRAFi-treated cells were plated at ∼2500 cells per well in a black, clear-bottom 96-well plate (Corning Inc., Corning, NY, USA). After cell seeding, RSL3, with or without Fer-1, was added the following morning, with media changes every three days (six replicates per condition). Plates were imaged using automated fluorescence microscopy (Cellavista Instrument, Synentec, Elmshorn, Germany). Twenty-five non-overlapping fluorescent images (20X objective, 5x5 montage) were taken twice daily for a total of 150 hours or until confluency. Cellavista image segmentation software (Synentec, Elmshorn, Germany) was utilized to calculate nuclear count (i.e., cell count) per well at each time point (Source = FITC, Dichro = FITC, Filter = FITC, Emission Time = 800 ms, Gain = 20x, Quality = High, Binning = 2x2). Cell nuclei counts across wells were normalized to the time of drug treatment and used to calculate drug-induced proliferation (DIP) rates, obtained as the slope of a linear fit to the log2-scaled population growth curve (94, 95). A dose-response curve was calculated across replicates using the ‘drm’ R package (96) with a 4-parameter log-logistic function, with DIP rate as the drug effect, as previously described (95). Replicates were used to calculate means and 95% confidence intervals for the dose-response curves. Data was visualized using the ‘ggplot2’ R package (version 3.2.0) (93).

## Model and experimental analysis code availability

The codes used to generate model simulations and analyze experimental data are publicly available at GitHub (github.com/SysBioCollab-UArk/Hayford_Melanoma_DTP_2026).

## Data availability

Sequencing data generated in this study have been deposited in the Gene Expression Omnibus (GEO) under accession GSE324655. Raw sequencing reads are available through the Sequence Read Archive (SRA) under BioProject accession PRJNA1431156. Processed data and analysis codes are available at GitHub (github.com/SysBioCollab-UArk/Hayford_Melanoma_DTP_2026).

## ACKNOWLEDGEMENTS

We thank Emily Hodges and Kelly Barnett for ATAC-seq reagents and experimental advice. We thank Jing Hao for reagent acquisition; Tony Capra, Christian Meyer, Sarah Maddox Groves, Carlos Lopez, Alissa Weaver, John McLean, Bruce Damon, and Ken Lau for useful discussions; and Rachana Nitin for providing feedback on an earlier version of this manuscript. B.B. thanks Nancy Cox and Christopher Wright for continued mentorship and support, and Jennifer (Piper) Below, Lea Davis, and Eric Gamazon for valued feedback and guidance. Sequencing studies were performed at the Vanderbilt Technologies for Advanced Genomics (VANTAGE) core facility at Vanderbilt University Medical Center with help from Angela Jones, Karen Beeri, Jamie Roberson, Latha Raju, and Matthew Scholz. Data processing and bioinformatics analyses were performed using computational resources at the Advanced Computing Center for Research and Education (ACCRE) at Vanderbilt University. We also acknowledge the Vanderbilt High Throughput Screening (HTS) core for contributions to this work.

This work was supported by the following funding sources: C.E.H., National Institutes of Health (NIH) Ruth L. Kirschstein National Research Service Award (NRSA, F31-CA221147) and Chemical-Biology Interface Training Grant (T32-GM065086); B.B., Integrated Biological Systems Training in Oncology Training Grant (T32-CA119925); P.E.S., Molecular Endocrinology Training Grant (T32-MH064913); B.B.P., Vanderbilt Institute for Clinical and Translational Research (VICTR) Grants (VR16721 and VR16721.1); A.B., NIH grants R01-CA255536 and U01-CA253540; V.Q., NIH Clinical and Translational Science Award (U54-CA113007); D.R.T., NIH Research Specialist Award (R50-CA243783); L.A.H., Vanderbilt Biomedical Informatics Training Program (NLM 5T15-LM007450-14), Quantitative Systems Biology Center at Vanderbilt, and National Cancer Institute (NCI) Transition Career Development Award (K22-CA237857). In addition, this material is based upon work supported by the National Science Foundation Graduate Research Fellowship Program under Grant No. 1937963 (to B.B). Any opinions, findings, and conclusions or recommendations expressed in this material are those of the authors and do not necessarily reflect the views of the National Science Foundation.

## AUTHOR CONTRIBUTIONS

Conceptualization: C.E.H., B.B., P.E.S., V.Q., D.R.T., L.A.H.; Experimental Data Collection: C.E.H., P.E.S., B.B.P., A.A., D.R.T.; Analysis and Interpretation of Results: C.E.H., B.B., P.E.S., V.Q., D.R.T., L.A.H.; Bioinformatics Analysis: C.E.H., B.B., P.E.S.; Writing: C.E.H., B.B., V.Q., L.A.H.; Review and Editing: C.E.H., B.B., P.E.S., A.B., V.Q., D.R.T., L.A.H.

## GLOSSARY

*This glossary defines key terms as used in this study, which may differ from standard usage in the literature*.

### Phenotype

The observable characteristics of a cell, such as morphology, migratory ability, and metastatic potential.

### Phenotypic state

A discrete category of closely related phenotypes that share common molecular or functional features. Individual cells are assigned to one phenotypic state at any given time. Although cells within a phenotypic state may vary at the molecular level, they are considered sufficiently similar to be classified together. Phenotypic states are also commonly referred to as “cell states” or “single-cell states.”

### Population state

A collection of cells distributed across a discrete set of phenotypic states. The population state refers to the equilibrium distribution of cells across these states under a defined set of conditions. At equilibrium, individual phenotypic states may be expanding or regressing, but their combined contributions determine the overall population behavior.

### DTP population state

A drug-induced population state in which the equilibrium distribution of cells across discrete phenotypic states results in a near-zero net proliferation rate for the population as a whole. Cells may transition among individual phenotypic states with differing proliferation rates under drug treatment, but their combined contributions balance so that the overall population size remains effectively constant.

---

**Figure S1: *Gene ontology analysis of single-cell transcriptomic clusters.* (A)** Ten single-cell clusters identified by the ‘Seurat’ software package used in this work (see Materials and Methods). In this UMAP projection, the “untreated small” (UTS) cluster corresponds to Seurat cluster 8, the “untreated large” (UTL) cluster to Seurat clusters 1, 2, 4, and 9, the “idling small” (IS) cluster to Seurat cluster 6, and the “idling large” (IL) cluster to Seurat clusters 0, 3, 5, and 7. **(B)** GO analysis of differentially expressed genes between the UTS and UTL clusters in the UMAP single-cell transcriptomics space. **(C)** Same as *B* but applied to the IS and IL clusters.

**Figure S2: *DNA barcode abundances and distributions across untreated and idling DTP states*. (A)** Heatmap of relative barcode abundances for all experimental replicates of the untreated and idling DTP populations. The heatmap is organized by decreasing barcode abundance in replicate 1 (R1) of the untreated condition. RPM: reads per million. (**B**) Number of unique barcodes in each treatment condition and number of shared barcodes between conditions. Lines correspond to the means of three experimental replicates. A minimum cutoff of 100 counts per million (CPM) was used. Note that the number of unique barcodes is higher in the drug-treated (idling) condition than in the untreated cells. We attribute this to biological noise, i.e., more lineages went extinct, by chance, in the untreated sample than in the idling sample. Overall, we interpret these data as evidence that lineages are not being clonally selected for upon drug treatment. **(C)** Proportional sharing of barcodes among experimental replicates for untreated and idling DTP populations. **(D)** Overlay of single cells from 20 barcoded lineages on the transcriptomic UMAP space from FIG. 1A. Colored contours (red and blue) reflect cell density for both treatment conditions.

**Figure S3: *Quality control of bulk ATAC-seq data and correlation analyses of transcriptome- and epigenome-based GO terms.*** (A) Unique reads versus sequenced reads for the bulk ATAC-seq data used in the epigenome analyses. **(B)** Insert size distribution of aligned reads from ATAC-seq data for untreated and idling DTP cells. Both conditions follow traditional nucleosome patterning. **(C)** Venn diagram of identified ATAC-seq peaks of open chromatin in the untreated and idling DTP populations. **(D)** Peak binding site distribution for untreated, idling DTP, and shared peaks. The x-axis represents the percentage of peaks; colors correspond to distances (kb) from the transcription start site (TSS). **(E)** Alignment of peaks to the TSS allows for the prediction of epigenomic features in untreated and idling DTP populations. **(F)** Correlation analysis between Biological Process (BP)-type GO terms from transcriptomic data (FIG. 3A) and epigenomic data (FIG. 3C). **(G)** Same as *F*, but for Cellular Component (CC)-type GO terms.

**Figure S4: *Ferroptosis pathway*.** Schematic diagram of the ferroptosis pathway from the WikiPathways database (https://wikipathways.org/pathways/WP4313.html) (81, 82). Colored boxes group genes associated with the GO terms listed at the bottom. Genes that are shaded are the ones included in the heatmap in FIG. 4A.

