## Supplementary Figures for "Heterogeneous, population-level drug-tolerant persisters exhibit ion-channel remodeling and ferroptosis susceptibility"

### Figure S1

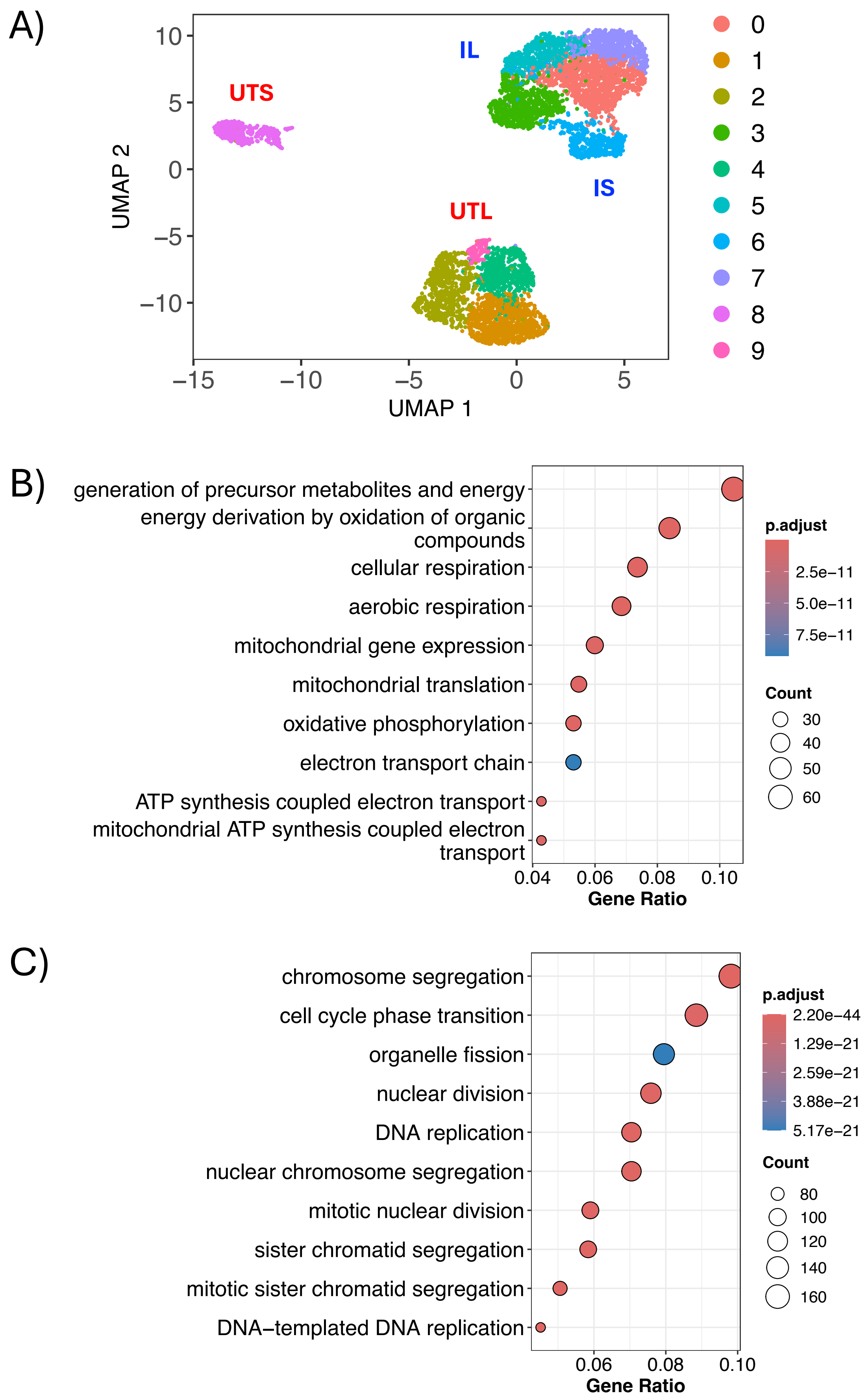
A)
UTS
IL
IS
UTL
UMAP 2
UMAP 1
10
5
0
−5
−10
−15
0
1
2
3
4
5
6
7
8
9
B)
generation of precursor metabolites and energy
energy derivation by oxidation of organic compounds
cellular respiration
aerobic respiration
mitochondrial gene expression
mitochondrial translation
oxidative phosphorylation
electron transport chain
ATP synthesis coupled electron transport
mitochondrial ATP synthesis coupled electron transport
0.04
0.06
0.08
0.10
Gene Ratio
p.adjust
2.5e−11
5.0e−11
7.5e−11
Count
30
40
50
60
C)
chromosome segregation
cell cycle phase transition
organelle fission
nuclear division
DNA replication
nuclear chromosome segregation
mitotic nuclear division
sister chromatid segregation
mitotic sister chromatid segregation
DNA−templated DNA replication
0.06
0.08
0.10
Gene Ratio
p.adjust
2.20e−44
1.29e−21
2.59e−21
3.88e−21
5.17e−21
Count
80
100
120
140
160

### Figure S2

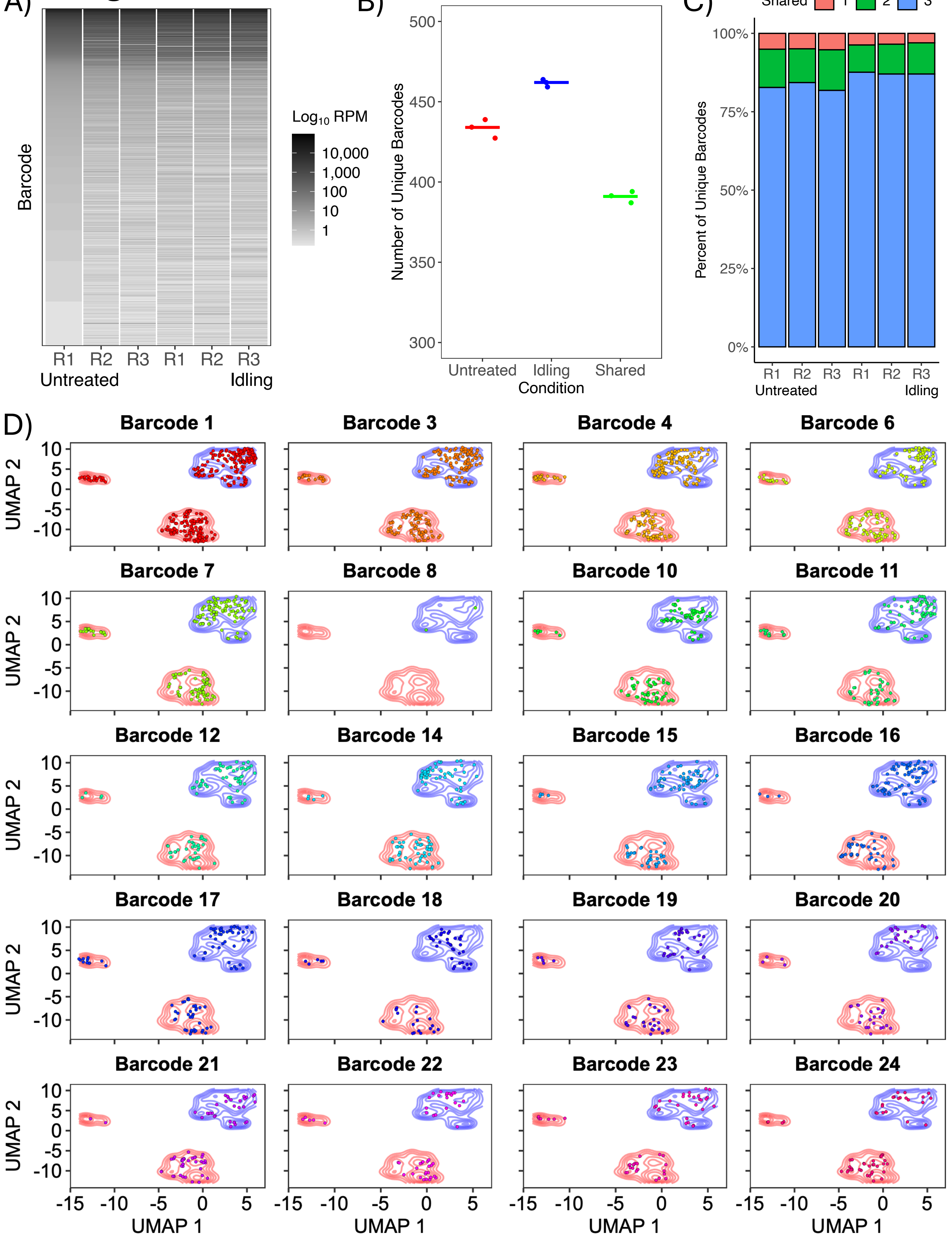
A)
Barcode
R1 R2 R3 R1 R2 R3
Untreated
Idling
Log10 RPM
10,000
1,000
100
10
1
B)
Number of Unique Barcodes
500
450
400
350
300
Untreated Idling Shared
Condition
C)
Shared 1 2 3
Percent of Unique Barcodes
100%
75%
50%
25%
0%
R1 R2 R3 R1 R2 R3
Untreated
Idling
D)
Barcode 1
Barcode 3
Barcode 4
Barcode 6
Barcode 7
Barcode 8
Barcode 10
Barcode 11
Barcode 12
Barcode 14
Barcode 15
Barcode 16
Barcode 17
Barcode 18
Barcode 19
Barcode 20
Barcode 21
Barcode 22
Barcode 23
Barcode 24
UMAP 2
UMAP 1
10 5 0 -5 -10
-15 -10 -5 0 5

### Figure S3

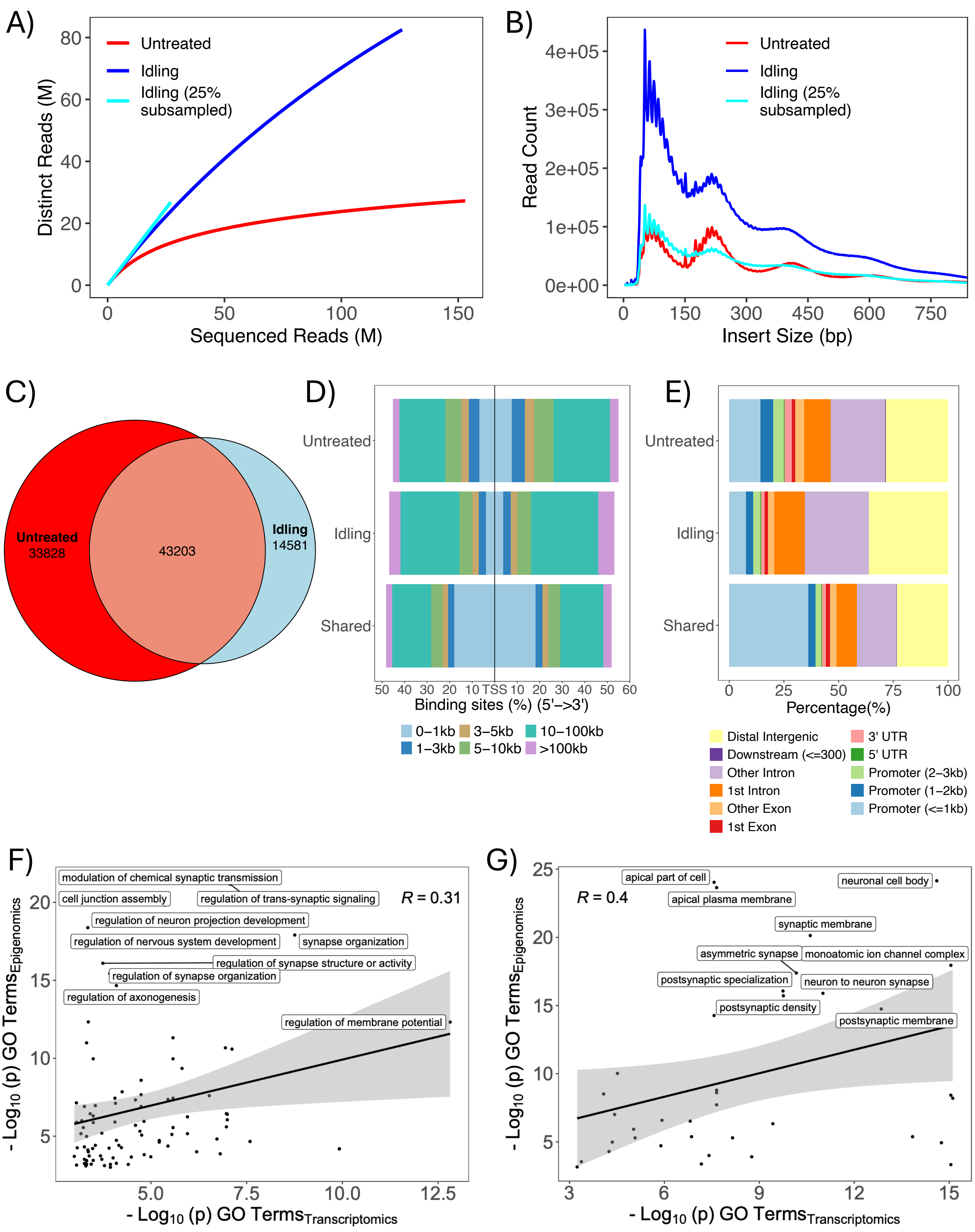
A)
Distinct Reads (M)
Sequenced Reads (M)
Untreated
Idling
Idling (25% subsampled)
B)
Read Count
Insert Size (bp)
Untreated
Idling
Idling (25% subsampled)
C)
Untreated 33828
43203
Idling 14581
D)
Untreated
Idling
Shared
Binding sites (%) (5'->3')
0–1kb
1–3kb
3–5kb
5–10kb
10–100kb
>100kb
E)
Untreated
Idling
Shared
Percentage(%)
Distal Intergenic
Downstream (<=300)
Other Intron
1st Intron
Other Exon
1st Exon
3' UTR
5' UTR
Promoter (2–3kb)
Promoter (1–2kb)
Promoter (<=1kb)
F)
modulation of chemical synaptic transmission
cell junction assembly
regulation of trans-synaptic signaling
R = 0.31
regulation of neuron projection development
regulation of nervous system development
synapse organization
regulation of synapse structure or activity
regulation of synapse organization
regulation of axonogenesis
regulation of membrane potential
- Log10 (p) GO Terms Epigenomics
- Log10 (p) GO Terms Transcriptomics
G)
apical part of cell
neuronal cell body
R = 0.4
apical plasma membrane
synaptic membrane
asymmetric synapse
monoatomic ion channel complex
postsynaptic specialization
neuron to neuron synapse
postsynaptic density
postsynaptic membrane
- Log10 (p) GO Terms Epigenomics
- Log10 (p) GO Terms Transcriptomics

### Figure S4

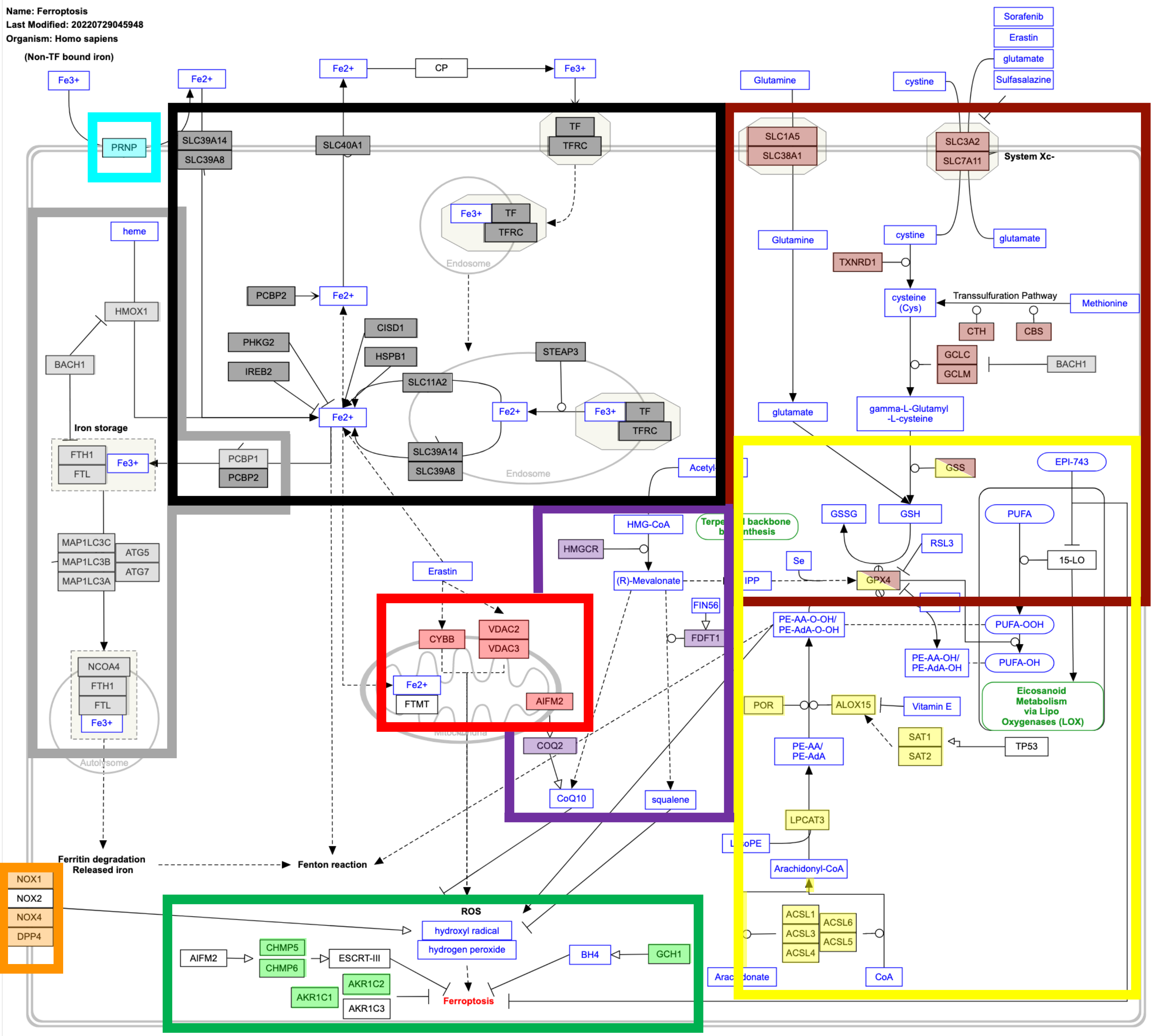


#### Gene ontology

- PUFA
- Terpenoid Biosynthesis
- Autolysosome Ferritin Storage
- Endosome Ferrous Transfer
- Mitochondria
- Superoxide Generation
- Inhibition of Ferroptosis
- Ferric acid reduction
- Glutathione Metabolism

wikipathways.org/pathways/WP4313.html
